# A multi-electrode array and immunofluorescence workflow to characterize impacts of HHV-6 infection in differentiated induced pluripotent stem cells

**DOI:** 10.64898/2026.09.22.753572

**Authors:** Elham Bahramian, Xue Yang, Ananya Bajpai, Jenifer Hernandez Garcia, Ruben Michael Ceballos

## Abstract

Roseoloviruses, notably human herpesviruses 6A and 6B (HHV-6A and HHV-6B), are neurotropic viruses implicated as agents in neurological disorders, including: epilepsy, multiple sclerosis, and chronic fatigue syndrome. However, the effects of roseolovirus infection on neuronal signaling and network activity are not characterized. This is, in part, due to the complexities of monitoring electrical activity in individual neurons and across neuronal connections during viral infection. This protocol describes a human induced pluripotent stem cell (iPSC)-derived neuronal culture system that employs multi-electrode array (MEA) recordings to study neurophysiological changes during viral infection. Two culture platforms are described: (a) NGN2-induced forebrain neuronal cultures; and, (b) progenitor cell-derived neuron–astrocyte mixed cultures. Cell composition in cultures is validated by immunofluorescence staining using neuronal, glial, and viral markers. Functional activity is assessed using extracellular recordings detected via the MEA2100 system. Mean firing rate, inter-spike interval, single-electrode bursting, and network burst activity are characterized between roseolovirus-infected versus uninfected (control) states. Pharmacological treatment with bicuculline, gabazine, and nicotine in conjunction with immunofluorescence is used to confirm functional responsiveness of defined neuronal neurotransmitter chemotypes. Here, we show that HHV-6A infection changes neuronal firing compared to uninfected controls. The methods/workflows described provide a strategy to study how virus infection alters neuronal excitability and may be adapted to compare the effects of different viruses on nerve cell function, infection time courses, different multiplicities of infection (MOI), and therapeutic interventions.

**SUMMARY:** Culturing nerve cells on embedded electrode array plates combined with parallel cultures on standard plates (and flasks) for immunofluorescence is an effective method for characterizing the impacts of virus infection on neuronal electrogenic function. Here we introduce a workflow to characterize the electrophysiological impacts of herpesvirus infection on cultured neurons.

## INTRODUCTION

Human herpesvirus 6 (HHV-6) is a neurotropic virus that is implicated in a range of neurological disorders^1–4^. Following infection of neurons and glial cells, roseoloviruses can trigger chronic neuroinflammation, gliosis, and disturbances in neurotransmitter systems, contributing to neuronal excitability and synaptic dysfunction.^4–6^ Although it is known that HHV6-A infects and establishes latency in the central nervous system (CNS)^7–9^, few studies have addressed the relative susceptibility and permissiveness of different nerve cell types.^5,7^ The relative infectivity and virulence of HHV-6A versus HHV-6B on different cell types also remains elusive. Electron microscopy studies confirm HHV-6B infection of oligodendrocytes.^10^ Antibody studies demonstrate that both HHV-6A and HHV-6B will bind antigens on oligodendrocytes.^11,12^ Glial fibrillary acidic protein (GFAP)-tagged astrocytes are shown to harbor HHV-6 antigens in vitro^13^ and in temporal lobe epilepsy patients.^5^ HHV-6A infection of astrocytes and Purkinje cells has been demonstrated both in vivo and in vitro.^14,15^ Recent studies from our laboratory using differentiated human neural stem cells (dHNSC) reveal that both HHV-6A and HHV-6B can infect glutamatergic and dopaminergic neurons but not GABAergic neurons in differentiated stem cells, suggesting a potential selective susceptibility of distinct neuronal neurotransmitter chemotypes.^13^ Our work also indicates differences in the magnitude and time-course of cytopathic effects (CPE) depending upon multiplicity of infection (MOI) as well as differences in relative virulence between HHV-6A versus HHV-6B on distinct nerve cell types.^13^ Such differences in cell tropism and relative virulence may underlie differences in symptomology and disease manifestation of the various neurological disorders reported to be associated with HHV-6 infection.

For example, in temporal lobe epilepsy (TLE) with mesial temporal sclerosis, HHV-6 DNA was detected in resected brain tissue from patients with drug-resistant epilepsy, indicating that inflammation induced by infection and neuronal injury may contribute to epileptogenesis.^16^ HHV-6 infection in limbic structures appears to promote gliotic scars and excitatory-inhibitory network imbalances, which also increase susceptibility to seizure.^17^ In multiple sclerosis (MS), HHV-6 infection has been implicated in both disease onset, persistence, and progression.^18^ Meta-analyses report significantly higher rates of HHV-6 detection in MS patients over controls, suggesting an etiologic role via mechanisms, such as molecular mimicry or immune dysregulation.^19^ Co-infection with Epstein–Barr virus (EBV) and HHV-6A has been shown to substantially increase the risk of MS development.^20^ In Alzheimer’s disease (AD), elevated levels of HHV-6 DNA have been detected in affected brain tissues compared to healthy controls, suggesting that viral-mediated neuroinflammation may exacerbate amyloid-β deposition and tau pathology.^21^ Polymicrobial interaction dynamics, including gut dysbiosis involving HHVs, are implicated in the neurodegenerative cascade and HHV translocation to the brain via vagal nerve cells.^22,23^ Chronic fatigue syndrome (CFS) and chronic HHV-6 infection have been associated with sustained neuroimmune activation, cognitive impairment, and fatigue.^24^ Quantitative studies confirm a high prevalence of HHV-6 infection in CFS patients compared to healthy individuals (i.e., controls).^24^ HHV-6 reactivation has also been proposed as a contributing factor in post-viral syndromes such as “long COVID”, with similar clinical features including chronic fatigue, mood disorders, and cognitive dysfunction.^25^

Viral infections can significantly disrupt neuronal electrophysiology, leading to impaired signaling and potential neurological disorders such as epilepsy. Human cytomegalovirus (HCMV) infection progressively reduces neuronal excitability by interfering with calcium signaling, ultimately silencing action potential generation in mature forebrain neurons.^26^ Similarly, COVID-19 has been linked to persistent EEG abnormalities such as reduced alpha rhythms, increases in slow waves (e.g., delta and theta waves), and epileptiform activity associated with neuroinflammation and neurovascular injury, resembling changes seen in neurodegenerative diseases like AD.^27^ Zika virus (ZIKV) initially increases neuronal activity but leads to noted loss of firing by 7 days post-infection, independent of inhibitory signaling, pointing to a progressive functional silencing of neurons.^28^ Enterovirus D-68 (EV-D68) also alters neuronal function, with microelectrode array data showing decreased spike and burst rates within hours of infection, suggesting disruption via viral replication or membrane changes rather than cytotoxicity.^29^

HHV-6 infection can significantly disrupt neuronal function by altering calcium signaling, neurotransmitter release, and synaptic transmission, key processes essential for maintaining neuronal excitability and thus neuronal network communication. HHV-6 has been shown to induce neuroinflammation, interfere with intracellular signaling pathways, and dysregulate calcium homeostasis, contributing to altered neuronal excitability, which is also linked to epilepsy and other neurological disorders. HHV-6 infection also affects glutamate release, potentially leading to excitotoxicity and further neuronal damage, thus impairing neural network signaling (i.e., electrophysiology) over the long term and increasing susceptibility to neurodegenerative conditions.^30,31^

The purpose of this study was to develop an effective multi-method workflow to characterize changes in electrophysiological signaling in response to viral challenge and map those changes to nerve cell chemotype. To achieve this goal, a multi-electrode array (MEA) plate system was employed to characterize electrophysiological perturbations in cultures of human induced pluripotent stem cell (iPSCs) differentiated into neurons and co-cultured with human astrocytes under uninfected versus HHV-6A infected conditions. The host-virus infection dynamics were monitored at different time points post-infection and compared to uninfected control cultures. 2-D and 3-D hydrogel (i.e., Matrigel®) cultures were tested along workflows consisting of both serial and parallel tracks to allow data integration across modes. Immunofluorescence using cell and virus antigen-antibody systems were used to identify cell types impacted by viral infection.

MEA platforms allow non-invasive, high throughput monitoring of extracellular electrical activity across cultured neuronal populations, capturing phenomena such as spike rate, burst behavior, and network connectivity.^32–34^ Unlike single-cell techniques (e.g., patch clamp), MEA enables parallel recordings from multiple regions of a culture simultaneously, providing an efficient way to monitor neuronal networks activity.^35,36^ Due to the ability to preserve cell viability while taking repeat measurements from the same culture over time, MEA systems are suitable for studying progressive infections (e.g., poly-viral infection) and long-term changes in neuronal activity.^37–39^ MEA studies have been extended from 2D monolayers to 3D neuronal assemblies, which more closely mimic in vivo environments and provide insights into how viral infections may alter functional synaptic integration and circuit dynamics in an intact organism.^32^

Advances in high-density MEA have further improved spatial resolution and sensitivity, allowing the detection of subcellular activity such as axonal transmission and propagation patterns. ^40,41^ Human iPSC-derived neurons were selected to study since they offer a physiologically-relevant in vitro model that recapitulates key molecular and electrophysiological properties of native neurons.^42^ Differentiated iPSCs develop functional synaptic networks and generate spontaneous and evoked action potentials, which is ideal for studying virus-induced alterations in neuronal excitability. Combined with immunofluorescence (IF) methods along both parallel and sequential workflows, key research questions may be addressed regarding the mechanisms by which viral infection alters nerve cell function. For example, neurons express cell surface receptors that are involved in viro-cell function such as virus entry and fast-acting (i.e., ion-gated) excitatory and inhibitory transmission. Therefore, IF data combined with MEA recordings can elucidate whether perturbations in individual neuron firing, or cell-to-cell transmission is due to dysregulation of receptor and/or ion channel expression and function.^43^

In this study, we utilized human iPSCs to generate neuronal cultures via two parallel workflows. Progenitor derived neuron-astrocyte cultures (Workflow A) and NGN2-induced forebrain neurons (Workflow B) were cultured on MEA plates and in a set of parallel plates (or flasks) for IF and other measures. Pharmacological tests using bicuculline, gabazine, and nicotine were performed as a validation step to verify that electrogenic systems in mature neuronal cultures are responsive prior to infection-based MEA analysis. We show the utility of combining MEA data with IF and other biochemical output to characterize the impacts of roseolovirus infection on nerve cells. Although it is beyond the scope of this paper, the workflows presented may also be combined with other methods (e.g., RT-qPCR) or -omics approaches (e.g., RNA-seq) to further elucidate molecular substrates underlying physiological outcomes from virus-host interactions.

## PROTOCOL

Ethics Statement: All work was performed in accordance with the guidelines and approval of the institutional Biosafety Committee (IBC) at University of California, Merced (IBC BUA No. R2803). The human induced pluripotent stem cell line and primary human astrocytes used in this protocol were originally derived from consented, de-identified donor material; their use does not constitute human subject research under 45 CFR 46 and did not require institutional Review Board approval. All procedures involving human herpesvirus 6 (HHV-6A/HHV-6B) were conducted under Biosafety Level 2 (BSL-2) containment following institutional biosafety guidelines and the approved IBC protocol.

### 1. Culture human induced pluripotent stem cells (hiPSCs)

Prepare two parallel cultures of human induced pluripotent stem cells (hiPSCs) as described for: (a) culturing on MEA plates to characterize electrophysiological responses and cytopathic effects; and, (b) culturing on multi-well plates (or in flasks) for biochemical and -omics analyses.

#### 1.1. Prepare Solubilized Basement Membrane Matrix (BMM)-coated cultureware

1.1.1. Thaw a stock 500 μL aliquot of undiluted BMM on ice (or at 4 °C overnight). Keep all reagents and pipettes pre-chilled to prevent gel formation.

1.1.2. Dilute the BMM at a 1:30 ratio with ice-cold DMEM/F-12 to prepare ∼15 mL of working matrix solution, which is sufficient to coat all wells of a multi-well plate (6- or 24-well plate) plus one MEA plate (see left most column Figure 1 for workflows A and B).

**Figure 1:**
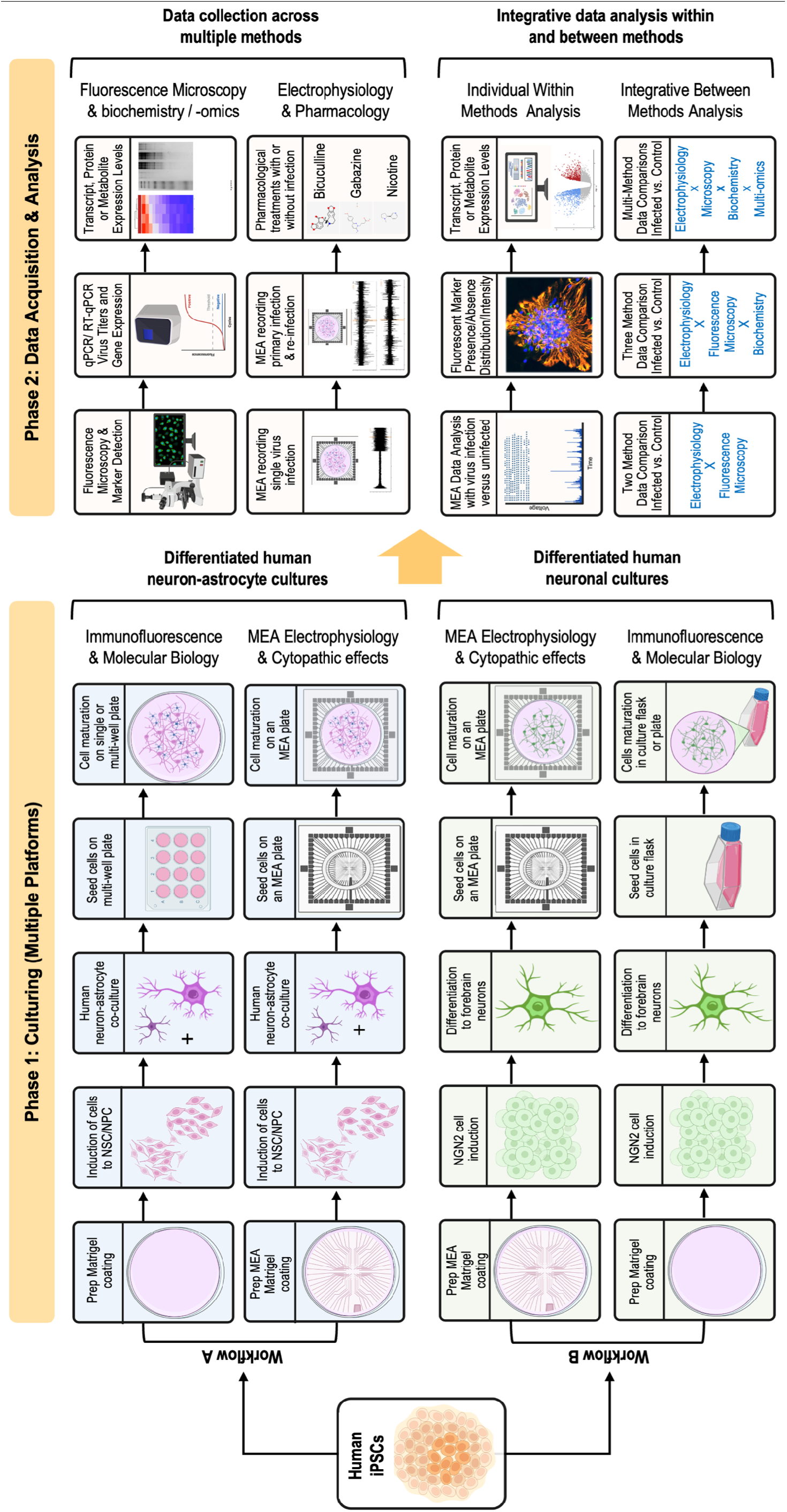
Workflow for MEA and multi-method analyses of virus-infected nerve cells. Culturing and differentiation of human iPSCs may be performed on MEA plates with parallel cultures in single or multi-well plates (or culture flasks). MEA plate cultures are used for electrophysiological analysis and parallel cultures may be used for complementary immunofluorescence microscopy, molecular biology, biochemistry, and/or multi-omics analyses (Phase 1). Predominantly neuronal cultures (e.g., NGN2-derived) or neuron-astrocytes cultures (e.g., NSC/NPC-derived) may be prepared in 2D (monolayer) or 3D (BMM/hydrogel) formats. Mature cultures may be subject to a host of experimental procedures, including: pharmacological assays (e.g., application of agonists/antagonists), molecular biological and genetics measures (e.g., qPCR, RT-qPCR), immunofluorescence, biochemical assays (e.g., PAGE, western blot), and multi-omics approaches (e.g., RNAseq), and others (Phase 2, top quadrant). Data from individual methods may be analyzed alone or integrated with other datasets to address complex questions in virology and virus ecology (Phase 2, bottom quadrant).

1.1.3. Dispense the diluted BMM onto the cultureware surface and swirl to ensure complete coverage. Incubate the coated cultureware at 37 °C for at least 1 h, then store sealed coated plates at 4 °C for up to 2 weeks, if needed.

#### 1.2. Thaw and seed hiPSCs

1.2.1. Pre-warm complete mTeSR-1 medium to room temperature and warm DMEM/F-12 containing 15 mM HEPES to 37 °C.

1.2.2. Thaw a cryovial of hiPSCs rapidly in a 37 °C water bath until only small ice crystals remain.

1.2.3. Transfer the thawed cells dropwise into 9 mL of pre-warmed DMEM/F-12 with 15 mM HEPES in a 15 mL conical tube.

1.2.4. Centrifuge the cell suspension at 300 x g for 5 min at room temperature, then aspirate the supernatant.

1.2.5. Resuspend the pellet in complete mTeSR-1 medium supplemented with 10 µM Y-27632.

1.2.6. Seed the cells onto matrix-coated cultureware at 9,000-18,000 cells/cm^2^ and incubate at 37 °C with carbogen (i.e., 95% O_2_/5% CO_2_).

1.2.7. Replace the medium the next day using matrix-coated medium without Y-27632.

NOTE: Supplier protocol (see Table of Materials) provides additional information regarding culturing of iPSCs.

#### 1.3. Passage hiPSCs

1.3.1. Inspect cultures daily for compact colonies with smooth borders and no obvious cell polarization.

1.3.2. Mechanically remove (e.g., with a sterile pipette tip or cell scraper) regions with any visibly differentiated cells before passaging.

1.3.3. Apply Accutase per the supplier protocol to passage colonies to new coated cultureware when they reach approximately 70-80% confluency, typically every 4-6 days.

NOTE: Y-27632 is a Rho-associated protein kinase (ROCK) inhibitor used transiently to improve cell survival following thawing, passaging, or single-cell dissociation. Add Y-27632 at 10 µM on the day of thawing, passaging, or seeding when cells are handled as single-cell suspensions. Remove Y-27632 at the next medium change, approximately 16–24 h later. Y-27632 is not included during routine maintenance or neuronal maturation unless cells undergo another single-cell dissociation step.

### 2. Prepare human astrocytes

Prepare cultures of human astrocytes as detailed below for co-culture with the NSC/NPC preparations described in section 1.0. Begin astrocyte expansion at least one week before the planned co-culture seeding day to ensure sufficient cells at passage.

#### 2.1. Prepare poly-D-lysine-coated cultureware

2.1.1. Prepare poly-D-lysine-coated cultureware before thawing the human astrocytes.

2.1.2. Use same procedure for coating cultureware as described in section 1.1.

#### 2.2. Thaw and seed human astrocytes

2.2.1. Thaw the cryopreserved human astrocyte vial rapidly in a 37 °C water bath until only small ice crystals remain.

2.2.2. Transfer the thawed cells directly into pre-warmed astrocyte maintenance medium in the poly-D-lysine-coated culture vessel.

2.2.3. Incubate at 37 °C with carbogen, undisturbed, for at least 16 h.

2.2.4. Replace the medium the next day to remove residual cryoprotectant and unattached cells.

#### 2.3. Maintain and passage human astrocytes

2.3.1. Replace the astrocyte maintenance medium every 3 days until the culture reaches approximately 70% confluency, then every other day thereafter.

2.3.2. Passage the astrocytes when the culture reaches approximately 90-95% confluency.

NOTE: Use astrocytes at passage 8 or lower for co-culture experiments. Avoid unnecessary freeze-thaw cycles. Information to retrieve the complete supplier’s protocol for thawing, culturing, and maintaining human astrocytes in vitro is provided in the Table of Materials (i.e., see the ScienCell protocol).

### 3. Prepare, clean, and sterilize MEA plates

Prepare MEA plates by oxygen plasma treatment followed by poly-D-lysine, and BMM coating. Use the cleaning and sterilization procedure described for coating and cell seeding. Do not autoclave MEA plates, as repeated autoclaving damages the insulation and electrode materials.

#### 3.1. Prepare clean and sterilize MEA plates

3.1.1. In a laminar-flow hood, gently add autoclaved distilled water along the inner wall of each MEA well with a sterile pipette until the recording surface is covered. Carefully remove the water from the edge of the well without touching the electrode field. Allow the plate to air-dry.

3.1.2. Immerse MEA plates in 1% Terg-A-Zyme solution at 4 °C overnight.

3.1.3. Carefully remove the Terg-A-Zyme solution and rinse the MEA recording surface with fresh autoclaved distilled water. Repeat as required to remove any remaining detergent residue.

3.1.4. Immerse the MEA plates in 70% ethanol for 15–20 min.

3.1.5. Rinse each MEA plate three times with fresh autoclaved distilled water, dispensing and removing water from the edge of the well without touching the electrode field. Allow the plates to air-dry briefly in the laminar-flow hood.

3.1.6. Expose the dried MEA plates to UV light in the laminar flow hood for 20–30 min.

#### 3.2. Oxygen plasma treatment of MEA plates

3.2.1. Treat the MEA plates with oxygen plasma to increase surface hydrophilicity and improve cell attachment.

3.2.2. Transfer each sterilized MEA plate into a labeled sterile 100 mm polystyrene Petri dish for handling and transport.

NOTE: This protocol uses standard 60-electrode MEA plates containing 59 recording electrodes and 1 internal reference electrode.

#### 3.3. MEA plate coating for cell seeding

3.3.1. On the day before seeding cells, add 100 µL of 50 µg/mL poly-D-lysine to the center of the MEA recording field and incubate at 4 °C overnight.

3.3.2. Rinse each MEA plate three times with autoclaved distilled water to remove residual poly-D-lysine.

3.3.3. Add 100 µL of 20 µg/mL laminin in DPBS to the MEA recording field and incubate at 37 °C for 1 h.

3.3.4. Aspirate the laminin solution without rinsing the surface.

3.3.5. Add 300 µL of 1:30 BMM to the MEA recording field and incubate at room temperature for 1 h before cell seeding.

NOTE: Use removable membrane chambers with O-rings during long-term culture. The membrane supports gas exchange while limiting evaporation and reducing contamination risk during extended neuronal maturation.

#### 3.4. Cell seeding on MEA plates

3.4.1. For monolayer “2D” Cell Culture Format, seed cells onto prepared MEA plates using the 1:30 dilution BMM coating over the electrode-embedded plate surface (also see section 4.0)

3.4.2. For gel Suspended “3D” Cell Culture Format, seed cells on MEA plates as a cell suspension consisting of a mixture of cells and a 1:1 BMM solution ∼300-500 μm thick (see section 5.0)

### 4. Generate human neuron cultures using two differentiation workflows

Monolayer 2D cultures as either neuron-astrocyte co-cultures (see Figure 1, Workflow A) or NGN2-induced forebrain neurons (see Figure 1, Workflow B) are developed according to the following procedures.

#### 4.1. Workflow A: NPC-derived neurons co-cultured with human astrocytes (2D format)

4.1.1. Maintain hiPSCs in complete mTeSR-1 medium on BMM-coated cultureware until healthy colonies are ready for neural induction.

4.1.2. Remove visibly differentiated regions from the hiPSC culture before neural induction.

4.1.3. Dissociate hiPSCs into a single-cell suspension with Accutase. Centrifuge cells at 300 x g for 5 min at room temperature.

4.1.4. Plate hiPSCs onto poly-D-lysine, laminin, and BMM-coated plates in complete neural induction medium with dual SMAD inhibition and 10 µM Y-27632. hiPSCs are induced into neural stem cells (NSCs) using NIM SMADi media.

NOTE: Add Y-27632 at 10 µM on the Day 0 in vitro whenever passaging or plating hiPSCs, NSCs, or neural progenitor cells (NPCs). See Table of Materials for coating material concentration.

4.1.5. Replace the medium daily with complete neural induction medium starting at Day 1 until cells are ready to passage.

4.1.6. Passage the cells with Accutase when neural rosettes have formed (typically Day 4-6).

4.1.7. Continue passaging neural stem cells approximately every 4-6 days. Maintain the cells in neural induction medium until the third passage.

4.1.8. Passage neural stem cells into complete neural progenitor medium at passage 3, approximately Day 18-21, and continue expansion until approximately Day 28 or until sufficient cells are available.

NOTE: In this protocol, the transition from neural stem cell (NSC) to neural progenitor cell (NPC) is defined operationally by the medium change at the third passage from NIM-SMADi to neural progenitor maintenance medium; no additional criterion is applied to distinguish the two states.

4.1.9. Cryopreserve neural progenitor cells at each passage for storage and batch consistency.

4.1.10. Confirm neural progenitor cell quality before downstream maturation by neural rosette morphology and, where possible, by immunostaining for PAX6, SOX1, and βIII-tubulin, together with absent or low expression of pluripotency markers.

4.1.11. Coat MEA plates sequentially with poly-D-lysine, laminin, and BMM as previously described (see section 3.3).

4.1.12. Dissociate neural progenitor cells and human astrocytes to single-cell suspensions in separate tubes using Accutase. Pellet each cell type at 300 x g for 5 min at room temperature.

4.1.13. Combine neural progenitor cells and astrocytes at a 4:1 ratio immediately before seeding in complete neural progenitor medium with Y-27632.

4.1.14. Seed the combined suspension onto the coated MEA plates at a total density of 80,000 cells per recording well. Dispense cells directly over the electrode field at the center of the well.

4.1.15. On Day 1 after seeding, replace the medium with complete neuronal maturation medium without Y-27632.

4.1.16. Replace half of the maturation medium every 2-3 days without disturbing the electrode field.

4.1.17. Mature the cultures for 38-50 days after seeding, or until stable spontaneous extracellular activity is detected by MEA recording.

NOTE: Workflow A uses hiPSCs that are first differentiated into neural stem cells then neural progenitor cells followed by maturation with human astrocytes before HHV-6 infection. Information to retrieve the complete supplier’s protocol for culturing astrocytes is provided in the Table of Materials.

#### 4.2. Workflow B: Generation of NGN2-induced forebrain neurons (2D format)

4.2.1. Dissociate hiPSC cultures into single cells and seed 50,000 cells per MEA onto MEA plates coated with poly-D-lysine and BMM. Seed the cells in complete seeding medium supplemented with 10 µM Y-27632 to support single-cell survival. Perform this initial seeding on Day −1 and allow the cells to attach overnight.

4.2.2. On the following day (Day 0), initiate neuronal differentiation by replacing the seeding medium with STEMdiff-TF Forebrain Induced Neuron Medium AB. Maintain a working volume of 0.5 mL per MEA.

4.2.3. Perform gentle medium changes according to the differentiation schedule provided by the supplier. Use Medium AB during the early induction phase, change to Medium B with PluriSIn-1 on Day 3, and then use Medium B alone on Day 4.

4.2.4. On Day 5, replace the differentiation medium with forebrain neuron maturation medium and maintain cultures until Day 21. Perform full medium changes every 2-3 days or half-medium changes every 2 days to reduce the risk of cell detachment. Supplement with uridine and FdU beginning on Day 7, as recommended by the manufacturer.

4.2.5. Maintain MEA cultures in a final working volume of 0.5 mL. During each medium change, aspirate and add medium slowly from the side of the MEA to avoid disturbing the coating matrix, neuronal layer, or electrode area.

NOTE: For continued maintenance after maturation, replace the maturation medium with complete BrainPhys medium and continue feeding cultures until infection and recordings.

NOTE: Workflow B uses hiPSCs converted into NGN2-induced forebrain neurons by lipid nanoparticle-mediated mRNA delivery before full maturation and HHV-6 infection. Human astrocytes are not added to Workflow B. Information to retrieve the complete supplier’s protocol for culturing NGN2-induced forebrain neurons is provided in the Table of Materials.

### 5. Plate cells on MEA plates for 3D Solubilized Basement Membrane Matrix (BMM) cultures

BMM-based 3D matrix cultures as neuron-astrocyte co-cultures (see Figure 1, Workflow A) or NGN2-induced forebrain neurons (see Figure 1, Workflow B) are developed according to the following procedures (also see Figure 1, two right-most columns).

#### 5.1. Prepare the cell-BMM suspension

5.1.1. Resuspend 60,000 neural progenitor cells plus 20,000 human astrocytes in 125 µL of complete BrainPhys differentiation neuronal medium.

5.1.2. Mix the cell suspension with cold BMM at a 1:1 ratio and keep the mixture on ice.

#### 5.2. Seed and solidify the 3D BMM culture on MEA plates

5.2.1. Apply the cell-BMM mixture onto the poly-D-lysine-coated center of each MEA, dispensing it as a confined dome over the electrode field so that the solidified gel forms a layer approximately 300–500 µm thick.

5.2.2. Incubate at 37 °C for 30 min to allow gel solidification.

5.2.3. Gently add 250 µL of BrainPhys neuronal medium over the solidified gel and incubate overnight at 37 °C with carbogen.

#### 5.3. Maintain the 3D BMM culture

5.3.1. On the following day, carefully remove the medium and replace it with fresh complete BrainPhys differentiation medium.

5.3.2. Perform medium changes every 2 days by carefully removing medium from around the gel and replacing it with fresh medium.

NOTE: Remove medium from the side of the well while tilting the MEA plate slightly. Do not touch the gel with the pipette tip.

### 6. Validate pharmacological responses of iPSC-derived neuronal cultures

Discerning the neurotransmitter chemotype of mature, differentiated neurons can be done several ways, including investigating neuronal responses to application of pharmacological agents as described below (also see Figure 1).

#### 6.1. Prepare pharmacological treatments

6.1.1. Prepare stock solutions of bicuculline, gabazine, and nicotine in sterile water at the concentrations specified in the Table of Materials.

6.1.2. Aliquot each stock solution and store at -20 °C protected from light.

NOTE: All three compounds in the salt or free-base forms used in this protocol are water soluble, and no dimethyl sulfoxide (DMSO) carrier is required.

6.1.3. On the day of the experiment, thaw a single-use aliquot and dilute it in pre-warmed neuronal maturation medium so that the final in-well concentration after addition is 10 µM.

#### 6.2. Record baseline and post-treatment MEA activity

6.2.1. Record baseline spontaneous activity from each MEA culture before treatment.

6.2.2. Add the working drug solution gently to the MEA well without disturbing the culture.

6.2.3. Record activity at the selected post-treatment time points, such as 2 h and 4 h after addition.

NOTE: Bicuculline, gabazine, and nicotine are pharmacologically active compounds. Handle all compounds according to institutional safety procedures. Bicuculline and gabazine are GABA-A receptor antagonists capable of inducing epileptic-like activity at the doses used.

### 7. Infect cultures with HHV-6 on MEA plates

Susceptibility of cultured neurons to viral infection may be assessed by monitoring neuron activity in response to viral challenge as described below (also Figure 1).

#### 7.1. Prepare cell-free HHV-6 stocks

7.1.1 Thaw frozen stocks of HHV-6A strain GS-infected cells and use them to infect uninfected HSB-2 host cells seeded at the required density (e.g., 1 × 10⁶ cells/mL) to generate cell-free virus.

7.1.2. Culture HHV-6-infected host cells in host growth medium for 5-7 days post-infection or until cytopathic effects are observed.

7.1.3. Collect cell-free culture supernatants, concentrate the virus with a 300 kDa molecular weight cutoff spin-concentration filter using high-speed centrifugation. Then, filter the retentate through a 0.45 µm sterile filter.

NOTE: Use appropriate institutional biosafety procedures and approved containment conditions for all risk-group 2 pathogen storage and handling.

#### 7.2. Quantify the number of viral genome copies by qPCR

7.2.1. Extract viral nucleic acid using a DNA extraction kit (see Table of Materials).

7.2.2. Elute DNA in 30 µL of nuclease-free water and store it at -20 °C.

7.2.3. Prepare a qPCR standard curve using known quantities of viral nucleic acid.

7.2.4. Prepare 20 µL qPCR reactions containing SYBR Green master mix, 0.5 µM each primer, and 2 µL of DNA sample or standard.

7.2.5. Use primers targeting the conserved HHV-6 U22 gene and perform a melting curve analysis to confirm specificity.

7.2.6. Use the standard curve to convert Ct values into viral genome copy number.

7.2.7. Adjust the copy number by sample input volume and extraction volume to report viral titer as genome copies per milliliter (GCN/μL).

#### 7.3. Infect neuronal cultures on MEA plates

7.3.1. Infect all cultures with cell-free HHV-6A or HHV-6B at an MOI of 1.

7.3.2. For 2D cultures, mix virus particles with 500 µL of differentiation medium and apply the inoculum directly to each MEA well using slow perfusion via a pipeter or peristaltic pump.

7.3.3. Incubate for 2 h at 37 °C, remove the inoculum, and replace it with 500 µL of fresh medium.

#### 7.4. Monitor infection dynamics versus uninfected control cultures

7.4.1. Monitor infected and uninfected control cultures at each post-infection time point using brightfield microscopy.

7.4.2. Compare cultures for visible cytopathic effects, including cell death, cell aggregation, syncytia formation, neurite loss, and detachment.

7.4.3. Record representative images for comparison with MEA recordings and qPCR results.

### 8. Perform immunofluorescent staining

Immunofluorescence imaging using an antibody-antigen system permits detection of cell types (e.g., neurons versus glia), neurotransmitter chemotype (e.g., glutamatergic vs. GABAergic cells), presence/absence and magnitude of expression of key cellular protein (e.g., surface receptors), or presence/absence and magnitude of expression of viral protein (e.g., envelope glycoproteins) as described below (also see Figure 1).

#### 8.1. Paraformaldehyde fixation of cells for immunofluorescence

8.1.1. In parallel with MEA cultures, grow companion cultures in 24-well plates. At each selected post-infection time point, aspirate the medium and rinse once with pre-warmed PBS.

8.1.2. Fix cells with 4% paraformaldehyde (PFA) in 1x PBS for 10 min at room temperature (RT).

CAUTION: PFA is hazardous; dispense in a fume hood and discard as chemical waste.

8.1.3. Wash 3 times for 5min each with 1x PBS.

NOTE: Fixed cultures may be held in PBS at 4°C for a few days, but stain viral antigens promptly, as HHV6 epitopes are prone to loss over storage.

#### 8.2. Permeabilization and blocking

8.2.1 Permeabilize with 0.1% Triton X-100 in 1x PBS for 10min at RT, then rinse once with 1x PBS.

8.2.2. Block with 1% bovine serum albumin (BSA) in PBS for 45 min at RT.

#### 8.3. Primary antibodies

8.3.1. Dilute primary antibodies from supplier stocks in blocking buffer to the working stock dilutions as follows: anti-βIII-tubulin (Tuj1/TUBB3, 1:500) for pan-neuronal identity; anti-VGLUT1 (1:500) for glutamatergic neurons; anti-tyrosine hydroxylase (TH, 1:500) for dopaminergic neurons; anti-GFAP (1:500) for astrocytes; and anti-HHV6 gp60/110 (1:300) for infected cells.

NOTE: The viral glycoprotein gp60/110 is a late/structural viral glycoprotein expressed only during lytic replication. This stain therefore labels cells undergoing active productive infection.

8.3.2. Co-stain no more than two markers per well, and pair primary antibodies raised in different host species so that species-specific secondaries do not cross-detect. Because TUJ1 and TH are both rabbit-hosted, do not place them in the same well.

8.3.3. Apply ∼300*μL* per well (24 well) and incubate overnight at 4°C in a humified chamber to prevent evaporation.

#### 8.4. Secondary antibodies

8.4.1. Wash each three times for 5 min with 1x PBS. Incubate with species matched, fluorophore conjugated secondary antibody (Typically 1:1000-1:2000) in blocking buffer for 1hr at RT, protected from light. Wash each, again, three times for 5min with 1x PBS.

8.4.2. Acquire uninfected (control) and infected conditions within the same imaging session using identical exposure and gain settings. Report these settings in the figure legend. Matched acquisition makes the absence of signal (over background) in uninfected controls interpretable.

#### 8.5. Nucleus counterstaining and imaging

8.5.1. Counterstain nuclei with 4’,6-Diamidino-2-phenylindole (DAPI) for 5 min, then wash once with 1x PBS.

8.5.2. Image well-plate cultures directly in 1x PBS. Acquire images on a fluorescence or confocal fluorescence microscope.

8.5.3. Expose the DAPI channel sufficiently to support downstream nuclear counting or segmentation.

### 9. Prepare parallel control flasks for viral titer quantification

Methods in quantitative polymerase chain reaction (e.g., qPCR or RT-qPCR) permit quantification of viral genomes as well as the quantification of cellular or viral transcripts (i.e., mRNA) during different post-infection points as described below (also see Figure 1).

#### 9.1. Seed and infect parallel control flasks

9.1.1. Seed iPSC-derived neuronal cells into tissue culture-treated flasks using the same differentiation medium and incubation conditions used for the 2D MEA cultures.

9.1.2. Use the same cell type, culture timing, and medium conditions as the matched MEA cultures.

9.1.3. Infect the flasks with HHV-6A at an MOI of 1 and maintain uninfected flasks as negative controls.

#### 9.2. Collect samples for viral DNA extraction

9.2.1. At time points corresponding to MEA recordings, collect one flask from each group.

9.2.2. Extract total DNA from each collected flask.

9.2.3. Store extracted DNA according to the DNA extraction kit instructions until qPCR analysis.

#### 9.3. Quantify viral titer by qPCR

9.3.1. Measure viral titer by qPCR^13^ and calculate HHV-6 genome copy number per μL (GC/μL) using a standard curve.

9.3.2. Report final titer as GCN/µL after adjusting for reaction volume and original sample input.

### 10. Record multi-electrode array activity

Electrophysiological recordings using an MEA system permit an assessment in neuronal activity changes as a function of: time-in-culture (no treatment; controls); application of pharmacological agents (or other chemicals); exposure to virus (or viruses); and, other experimental variables as described below (also see Figure 1).

#### 10.1. Prepare the recording system

10.1.1. Power on the MEA2100-mini headstage and temperature controller. Set the temperature controller to 37 °C and allow the headstage to equilibrate.

10.1.2. Launch the data acquisition software and load the recording configuration file. Set the sampling rate to 10–25 kHz and apply a 200–3000 Hz bandpass filter to isolate spike-band activity.

10.1.3. Maintain the recording temperature at 37 °C using an incubator-compatible setup or external temperature control unit. If using temperature control, set the temperature controller to 37 °C and allow the headstage to equilibrate.

NOTE: Acquire all recordings with the data acquisition software according to the manufacturer’s manual. Use the same recording configuration for all cultures and time points.

#### 10.2. Mount and verify the MEA plate

10.2.1. Transfer the 60MEA200/30iR-Ti-gr MEA plate from the incubator and mount it on the MEA2100-mini headstage. Insert the plate securely and check electrical contact in the software.

10.2.2. Select all recording channels and verify that signal paths are mapped to the physical electrodes.

10.2.3. Allow the MEA plate to equilibrate on the headstage for 5 min before acquisition.

#### 10.3. Record and save MEA activity

10.3.1. Record spontaneous extracellular activity for 5 min per MEA culture at each time point.

10.3.2. Stop the recording, save the file for offline analysis, and return the MEA plate to the incubator immediately after recording.

NOTE: Minimize the time each plate spends outside the incubator. When recording multiple plates, schedule sequential recordings so that each plate remains outside the incubator for the shortest possible time.

### 11. Analyze microelectrode array data

Electrophysiological recordings from MEA raw data provide useful measures, including: mean firing rate; inter-spike interval time; burst rate; and, other measures. These data may be compiled and presented as described below (also see Figures 2, 5, and 6).

**Figure 2:**
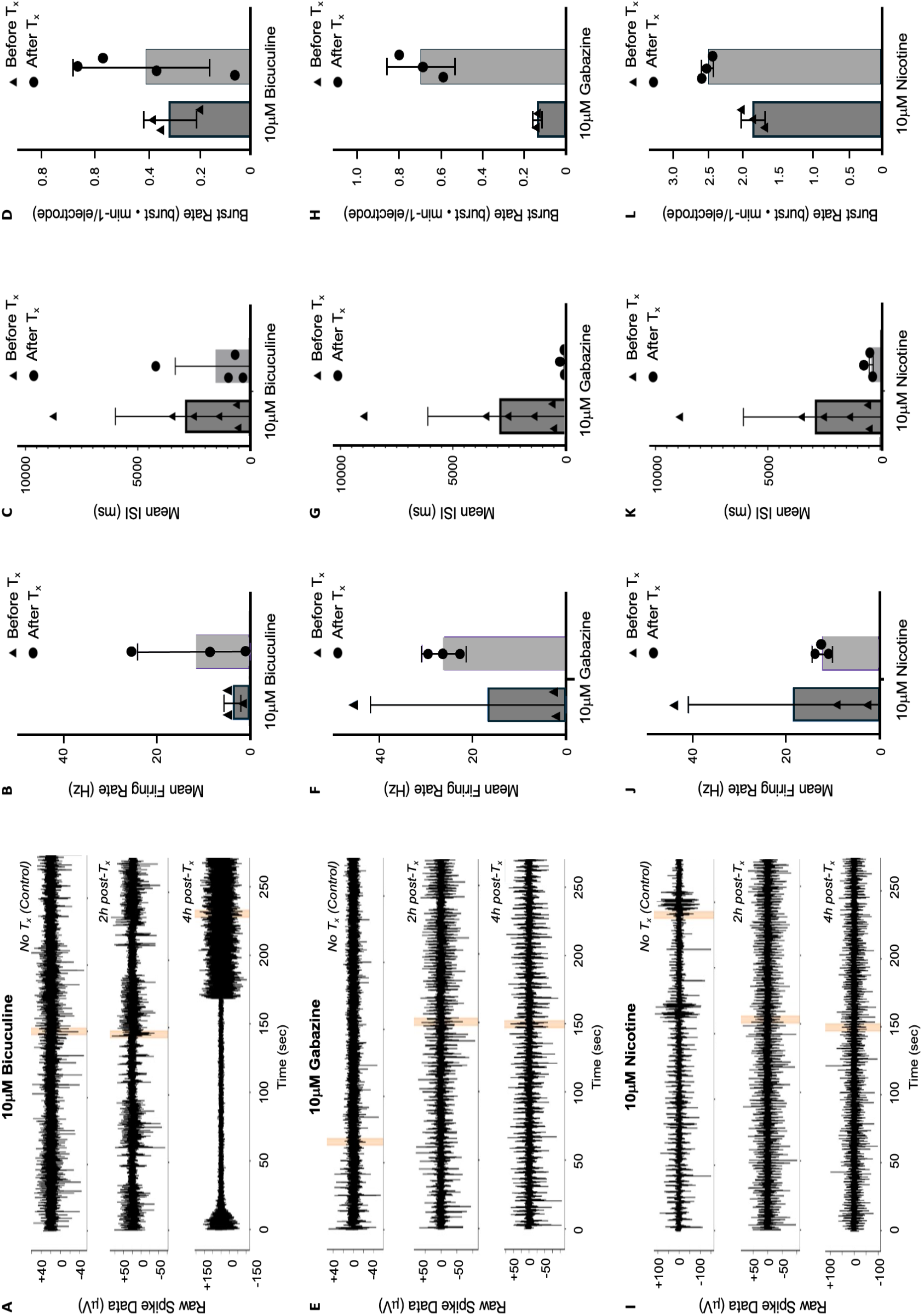
Pharmacological validation of progenitor cell-derived neuron–astrocyte cultures. Raw spike data from MEA recordings before treatment, 2 h after treatment, and 4 h after treatment with: **(A)** 10 µM bicuculline; **(E)** 10 µM gabazine; and, **(I)** 10 µM nicotine. Recording time window analyzed (orange highlighted regions). Mean Firing Rate (MFR) before and after HHV-6A infection at MOI = 1 for: **(B)** bicuculline-treated; **(F)** gabazine-treated, and, **(J)** nicotine-treated cultures. Mean inter-spike interval (ISI) before and after treatment with: **(C)** bicuculline; **(G)** gabazine; and, **(K)** nicotine. Burst rate per electrode (BRE) before and after treatment with: **(D)** bicuculline; **(H)** gabazine; and, **(L)** nicotine.

#### 11.1. Export time-stamped MEA data

11.1.1. Export time-stamped MEA data for recordings as an HDF5 file via DataManager software.

11.1.2. .HDF5 files from MEA recordings (each file = one sample = one condition and time point).

11.1.3. Content: Each file contains multiple electrodes with a list of spike timestamps.

11.1.4. Use Python for MEA data analysis. Use h5py to open HDF5 recording files. Use NumPy to perform numerical calculations on spike timestamps and electrode-level data. Use Matplotlib to generate summary plots of firing rate, inter-spike interval, burst rate, and network activity. Use pandas to organize results into tables by sample, condition, time point, and electrode.^44–46^

#### 11.2. Extract and map spike timestamp data

11.2.1. Extract spike timestamp datasets from the SegmentData_ts_* fields and calculate recording duration from the analog data stream sampled at 10 kHz.

11.2.2. Convert spike timestamps to seconds according to the units stored in the HDF5 file.

11.2.3. Map spike data to the 60 physical MEA channels. Exclude extra unmapped SegmentData_ts_* datasets if more than 60 timestamp datasets are present.

#### 11.3. Calculate electrode-level activity metrics

11.3.1. Define active electrodes as physical electrodes with at least 10 spikes during the full 5-min recording. Calculate active electrode number and percentage for each file.

11.3.2. Calculate mean firing rate (MFR) for each active electrode as total spike count divided by recording duration in seconds. Report MFR across active electrodes for each recording.

11.3.3. Calculate inter-spike intervals (ISI) from electrodes with at least 20 spikes by sorting timestamps and calculating the time differences between consecutive spikes. Exclude silent intervals longer than 10 s and report the median ISI for each recording.

#### 11.4. Detect single-electrode and network bursts

11.4.1. Detect single-electrode bursts using an ISI-based method. Define a burst as at least five spikes with consecutive ISIs of 100 ms or less and a burst duration between 20 ms and 2 s.

11.4.2. Calculate burst rate as the number of bursts divided by recording duration in minutes. Apply the same exclusion threshold across recordings to reduce distortion from isolated hyper-bursting channels.

11.4.3. Detect network bursts by combining spike timestamps from all active electrodes and binning spikes into 100 ms windows. Define candidate network bursts as bins exceeding the recording-wide population threshold and require participation of at least 20% of active electrodes and at least five electrodes.

11.4.4. Merge consecutive positive bins into one network burst event and calculate network burst rate as network bursts per minute.

### 12. Statistical analysis

Statistical summaries of MEA-derived electrophysiological parameters are generated from the electrode-level measurements obtained during each 5-minute recording as described below.

#### 12.1. Define experimental and technical measures

12.1.1 Treat each MEA recording file as one experimental sample representing a defined culture condition and recording time point.

12.1.2 Consider each individual active electrode within each MEA run as technical measurement of electrical activity within that culture (as opposed to an individual biological replicate).

12.1.3 Define the number of active electrodes contributing to each recording as described in section 11.3.1. The number of active electrodes is reported for the corresponding infection time points shown in the electrophysiological traces.

#### 12.2. Summarize electrophysiological data

12.2.1. Summarize mean firing rate (MFR) and mean burst rate (MBR) across active electrodes within each recording. Calculate and report median inter-spike interval (ISI) as described in section 11.3.3.

12.2.2. Display electrode-level variability using error bars where applicable and show individual measurements as available (see Figure 2).

#### 12.3. Interpret comparisons between experimental conditions

12.3.1 Longitudinal MEA recordings represent repeated measurements of the same cultures, and the number of independent biological replicates is limited. Therefore, interpret differences between recording time points descriptively (i.e., greater than v. less than, increase v. decrease). Only in cases where multiple true biological replicates are available in with an N > 8, which would represent months of work with only one MEA system, do not consider individual electrodes as independent biological replicates for inferential statistical modeling.

12.3.2 Describe increases and decreases in electrophysiological parameters as observed changes in recorded cultures (e.g., as a function of time and treatment). Do not infer statistical significance from electrode-level variation alone.

#### 13. Integrative analyses

Results from the electrophysiological, immunofluorescence, biochemical, and -omics datasets may be integrated to address research questions.

#### 13.1. Compare MEA activity with -omics data

13.1.1 Determine if there are correlations between electrophysiology (e.g., mean firing rate) and viral genome copy number (or changes in expression of cellular or viral genes of interest) as a function of time in virus-infected versus uninfected (control) cultures.

13.1.2 Determine if there are correlations between cytopathic effects (e.g., neurite extension) and electrophysiological data (network burst rate) aligned with questions of scientific interest in virus-infected and uninfected (control) cultures.

#### 13.2. Compare MEA activity with immunofluorescence and biochemical data

13.2.1. Determine if there are correlations between electrophysiology (e.g., ISI, burst rate) immunofluorescence data (e.g., neuronal cell types present) for virus-exposed versus uninfected (control) cultures.

13.2.2. Integrate MEA results with immunofluorescence, qPCR, transcriptomic, or other molecular analyses to identify pathways that may contribute to altered neuronal firing, bursting, and network activity, including any other cellular or viral biochemical markers (e.g., receptors) in virus-exposed versus uninfected control cultures.

## RESULTS

This protocol demonstrates that parallel cultures of differentiated neurons may be generated using distinct differentiation protocols for both neuronal and astrocyte-neuron mixed cultures. Furthermore, cells may be cultured in multi-well plates, petri-style culture plates, culture flasks, or MEA plates. Both 2D monolayer and 3D cell suspension BMM (i.e., Matrigel) result in vial cell growth (Figure 1, Phase 1). Once cells are fully differentiated into electrogenic neurons, multiple techniques may be used to collect different types of data (Figure 1, Phase 2, top) which when analyzed individually and in concert, provide insights into the impacts of neurotropic virus infection on cell signaling (Figure 1, Phase 2, bottom).

### Pharmacology and Electrophysiological Responses

To determine if neurons are electrogenic and to determine which neuronal neurotransmitter chemotypes may be active in culture, pharmacological agents may be employed (Figure 2). In this study NSC/NPC-derived neuron and astrocyte co-cultures were subjected to 10μm bicuculine, a potent antagonist of GABA_A_ receptors (Figure 2, panels A-D). After treatment (T_x_) with bicuculine, it appears that mean firing rate (MFR) increases in the culture and inter-spike interval (ISI) drops, which would be consistent with system excitation. This is further supported by an observable increase in burst rate after treatment. Visual inspection of the raw spike data, certainly at 2 hr post-T_x_ (Figure 2, panel A, middle) clearly demonstrates the ability of bicuculine to induce burst activity compared to the untreated control (Figure 2, panel A, top), suggesting the presence of GABAergic neurons in the mixed culture. In a separate trial, NSC/BNPC-derived neuron-astrocyte co-cultures were also treated with gabazine, another GABA_A_ receptor antagonist. Again, MFR appears to increase, and ISI appears to decrease but one-off outliers result in no statistical significance (Figure 2, panels F and G, respectively). However, burst rate does show a notable increase in these 10 μm gabazine treated cultures (Figure 2, panel H) providing confidence that GABAergic cells are active in culture. In yet another trial, NSC/NPC-derived neuron-astrocyte co-cultures were treated with 10μm nicotine, a strong cholinergic agonist that acts on a myriad of fast-acting nicotine acetylcholine receptors. MFR does not appear to significantly change with the application of nicotine. Indeed, there may even be a decrease in MFR; however, there is an apparent drop in ISI. Again, due to outliers, no statistical significance emerges from MFR and ISI data (Figure 2, panels J and K). However, there is a readily observable increase in burst rate after application of nicotine (Figure 2, panel L), suggesting that there may be cholinergic cells within the culture.

### Immunofluorescence and Brightfield Microscopy

To further explore not only the presence or absence of neuronal neurotransmitter chemotypes but also the ability of HHV-6A to infect select nerve cell types, immunofluorescence (IF) was used. For NSC/NPC-derived neuron-astrocyte co-cultures, parallel cultures were exposed to HHV-6A at MOI = 1 at day 51, each targeting HHV-6A gp60/110 and a select antigen with a fluorescent antibody (Figure 3).

**Figure 3:**
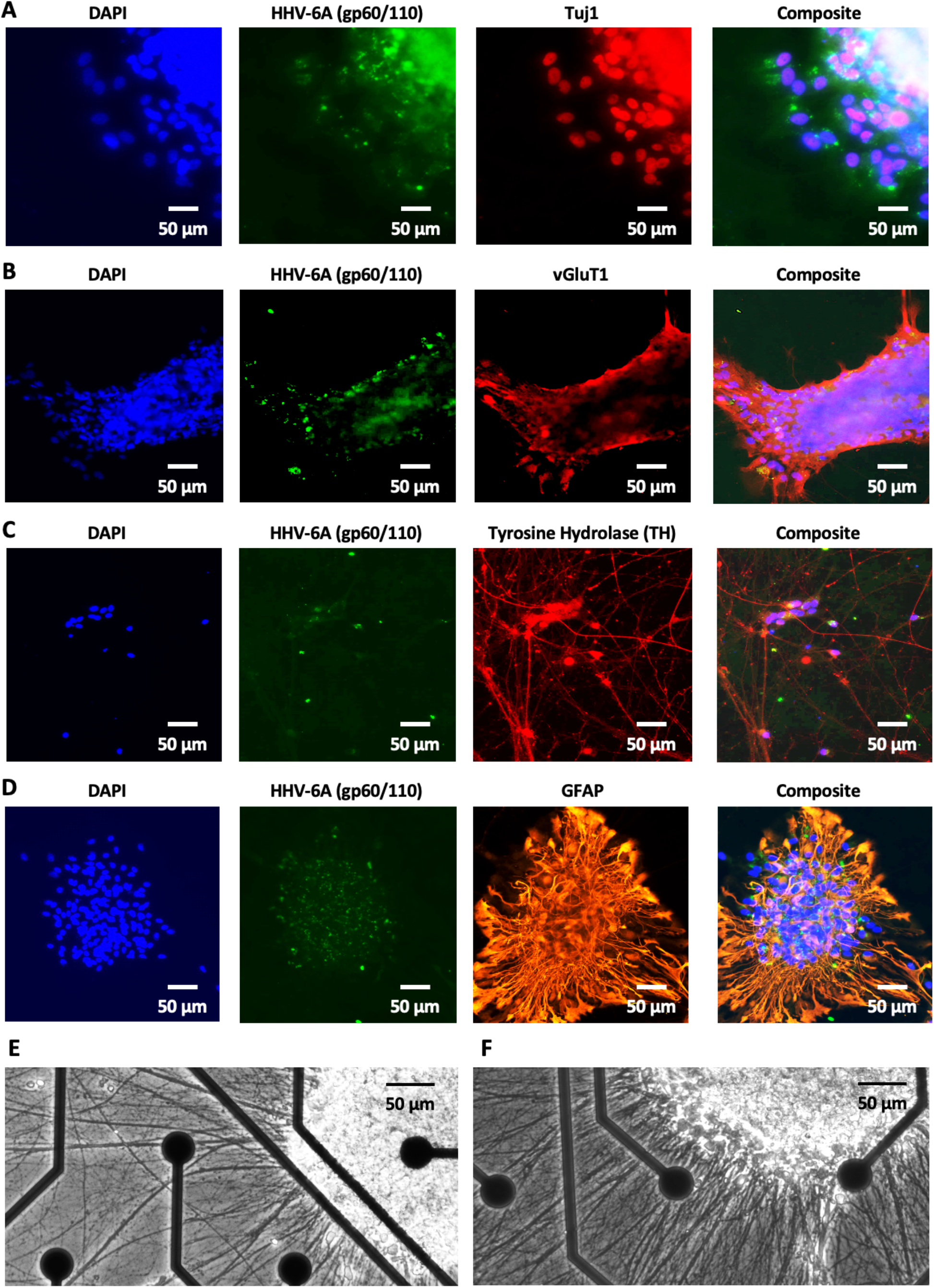
Immunofluorescence and brightfield imaging of neuron–astrocyte cultures. Parallel cultures of NSC/NPC-derived neuron-astrocyte cultures were infected with HHV-6A at MOI = 1. For all cultures, fluorescent antibodies targeting the gp60/110 envelope glycoprotein of HHV-6 (green) and DAPI to identify cell nuclei (blue) were used. (A) A fluorescent antibody targeting βIII-tubulin (e.g., Tuj1), a neuron-specific antigen (red), was used to assess whether HHV-6A infects neurons as indicated by overlapping fluorescence emissions (red x blue x green). (B) A fluorescent antibody targeting vGluT1, a glutamatergic cell-specific antigen (red), was used to assess whether HHV-6A infects glutamatergic neurons as suggested by overlapping fluorescence. (C) A fluorescent antibody targeting tyrosine hydrolase (TH), a catecholaminergic cell-specific antigen (red), was used to assess whether HHV-6A neurons catecholaminergic neurons (e.g., dopaminergic cells) as would be suggested by overlapping fluorescence signals. (D) A fluorescent antibody targeting GFAP, a glia-specific antigen (orange), was used to assess whether HHV-6A infects astrocytes as indicated by overlapping signals (orange x blue x green). Brightfield microscopy reveals cell aggregation (with no apparent syncytia), robust cell soma, and extensive neurite extensions/networks in MEA cultures (E, F). Scale bar: 50 μm.

IF data exhibit notable overlap in DAPI, HHV-6A gp60/110, and βIII-tubulin (i.e., Tuj1) fluorescence (Figure 3, row A), indicating that neurons are susceptible and permissive to HHV-6A infection. Likewise, IF data show overlap between the HHV-6A gp60/110 fluoroprobe and fluorescence from the antibody targeting vGluT1, an antigen used to detect the presence of glutamatergic neurons (Figure 3, row B), suggesting that glutamatergic cells are also susceptible. To assess the susceptibility of catecholaminergic cells (e.g., dopamine neurons) to HHV-6A infection, a fluorescent antibody-antigen system was used to detect the presence of tyrosine hydrolase (TH) in these NSC/NPC-derived neuron-astrocyte co-cultures (Figure 3, row C). Although HHV-6A gp60/110 signals are sparse, there is some notable overlap with TH targeted fluorescence and DAPI signal, suggesting that catecholaminergic cells may also be susceptible to HHV-6A infection. To test the susceptibility of glia to HHV-6A, glial fibrillary acidic protein (GFAP), a marker highlighting the presence of astrocytes, was used (Figure 3, row D). HHV-6A gp60/110 antibody fluorescence clearly overlaps with both DAPI and GFAP antibody fluorescence. Brightfield microscopy reveals mature neurons with expansive neurite networks and healthy cells (Figure 3, panels E and F). To assess the consistency of these results with NGN2-derived neurons, a similar set of trials were conducted.

For NGN2-derived neurons, a set of parallel culture were exposed to HHV-6A at an MOI = 1 on day 67 of the culture (Figure 4). IF images exhibit a notable overlap between HHV-6A gp60/110 and βIII-tubulin (i.e., Tuj1) targeted antibody fluorescence as well as with DAPI fluorescence (Figure 4, row A), indicating that HHV-6A productively infects neurons from the NGN2 cultures. IF images likewise show that HHV-6A gp60/110 signal overlaps with a cell clump that is positive for vGluT1 (Figure 4, row B). Although HHV-6A gp60/110 signals are sparse, there is some notable overlap with TH targeted fluorescence and DAPI signal, suggesting that catecholaminergic cells are also susceptible to HHV-6A infection in NGN2-derived neurons (Figure 4, row C). Brightfield microscopy of NGN2-derived cells show cellular aggregation into clumps and neurite networks (Figure 4, panels D and E), similar to that found in NSC/NPC-derived neuron-astrocyte cultures.

**Figure 4:**
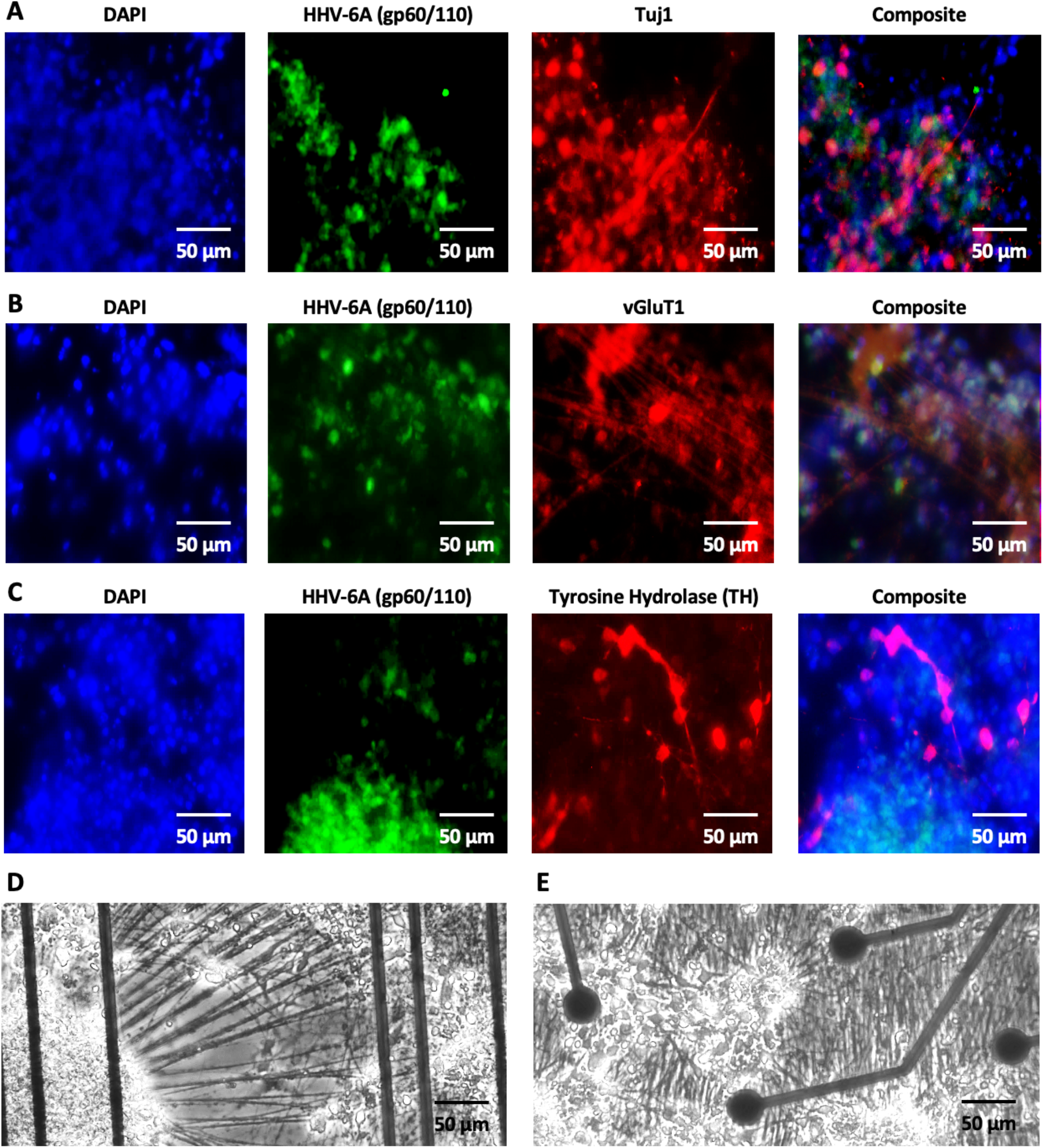
Immunofluorescence and brightfield imaging of NGN2-derived forebrain neurons. Parallel cultures of NGN2-inducible forebrain neurons were infected with HHV-6A at MOI = 1. For all cultures, fluorescent antibodies targeting the gp60/110 envelope glycoprotein of HHV-6 (green) and DAPI to identify cell nuclei (blue) were used. (A) A fluorescent antibody targeting βIII-tubulin (e.g., Tuj1) was used (red) to assess whether HHV-6A infects neurons as indicated by overlapping fluorescence emissions (red x blue x green). (B) A fluorescent antibody targeting vGluT1 (red) was used to assess whether HHV-6A infects glutamatergic neurons as suggested by overlapping fluorescence. (C) A fluorescent antibody targeting TH (red) was used to assess whether HHV-6A neurons catecholaminergic neurons (e.g., dopaminergic cells) as would be suggested by overlapping fluorescence signals. Brightfield microscopy reveals cell aggregation (i.e., clumps), robust cell soma, and extensive neurite extensions/networks in MEA cultures (D, E). Scale bar: 50 μm.

### Electrophysiology and Electrogenic Cell Responses to Viral Infection

MEA recordings provide a functional readout of changes in neuronal activity in response to infection by HHV-6A in both neuron-astrocyte co-cultures and NGN2-induced neuronal cultures. From each 5-min spontaneous recording, mean firing rate, median inter-spike interval (ISI), single-electrode burst rate, and the number of active electrodes were monitored under uninfected (control) conditions and for multiple time-points after infection with HHV-6A at an MOI=1 for differentiated nerve cells in NSC/NPC-derived neuron-astrocyte cultures (Figure 5).

**Figure 5:**
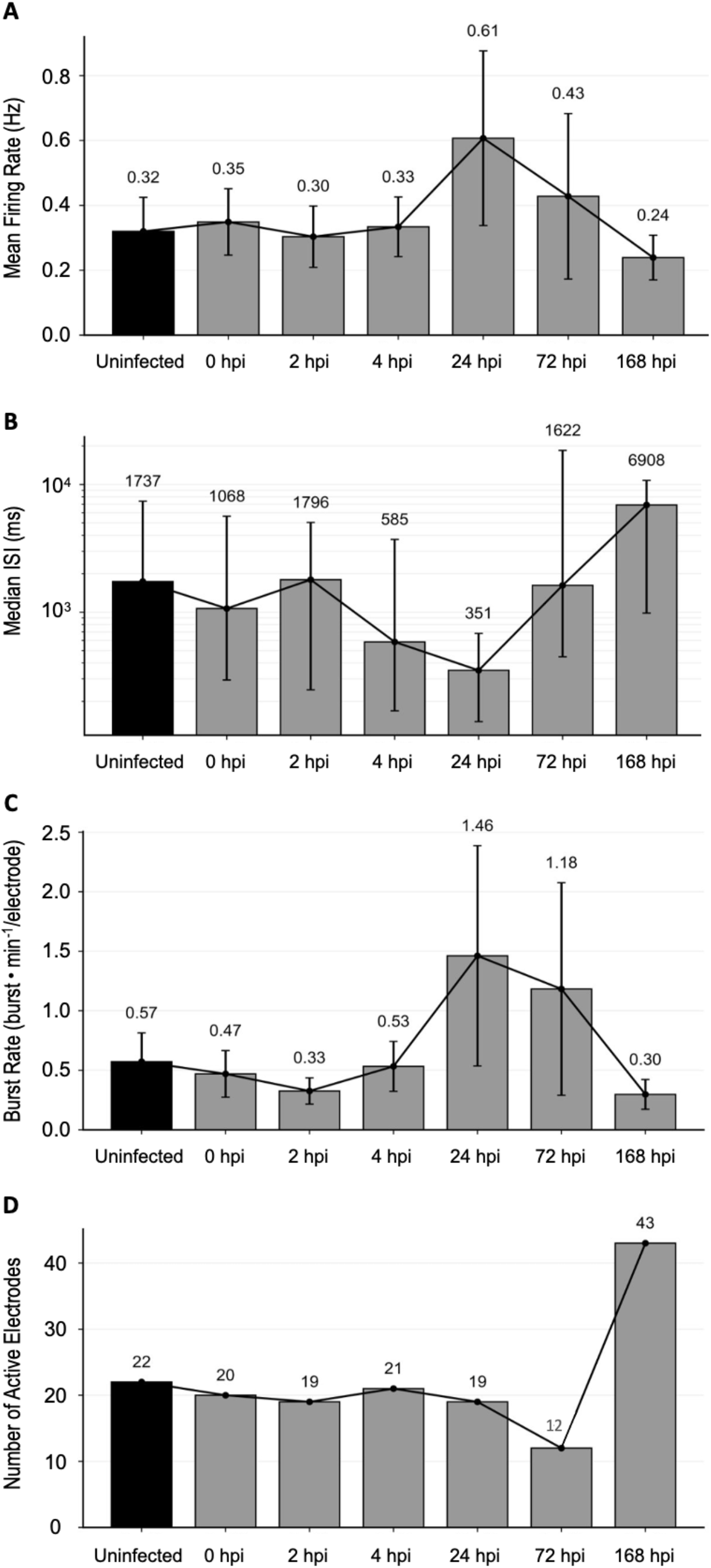
NSC/NPC-derived neuron-astrocyte electrophysiology upon HHV-6A infection. MEA recordings were used to assess electrophysiological activity in progenitor cell-derived neuron-astrocyte cultures at 51 days in culture both before and after HHV-6A infection (MOI=1). **(A)** Mean firing rate (MFR) across active electrodes. **(B)** Median inter-spike interval (ISI). **(C)** Burst rate per active electrode (BRE). **(D)** Number of active electrodes. Bars show summary values across active electrodes with error bars representing variability across active electrodes where applicable; the black line connects time points to show the longitudinal trend. Black bars indicate the uninfected baseline, and gray bars indicate HHV-6A infection time points.

Mean firing rate remains steady during the initial hours after infection (0-4 hpi) with a noted increase in firing at 24 hpi (Figure 5, panel A) with a steady decline across later recording times (72 and 168 hpi). Inter-spike interval (ISI) inversely maps to changes in firing rate with a notable dip at 24 hpi (Figure 5, panel B) and subsequent increase across later time-point (72 and 168 hpi). Burst rate likewise maps with a positive correlation to mean firing rate with a notable increase in firing at 24 hpi (Figure 5, panel C) and a gradual decline across latter time-points (72 and 168 hpi). No significant losses in electrode activity are observed during recording however, after 7 days post-infection, it appears that additional electrodes are engaged (Figure 5, panel D). Consistent electrode activity (i.e., number of active electrodes engaged) provides confidence that changes in electrical responses are due to the impacts of viral infection rather than the loss (or gain) of cell activity for these NSC/NPC-derived neuron-astrocyte co-cultures examined by MEA after 51 days in culture.

For NGN2-derived neuronal cultures, mean firing rate, median ISI, and burst rate measures were also collected but twice – once at 29 days in culture (i.e., primary infection) and, again, as a reinfection at 67 days in culture (i.e., secondary infection) -with MEA recordings taken during each time period (Figure 6).

**Figure 6:**
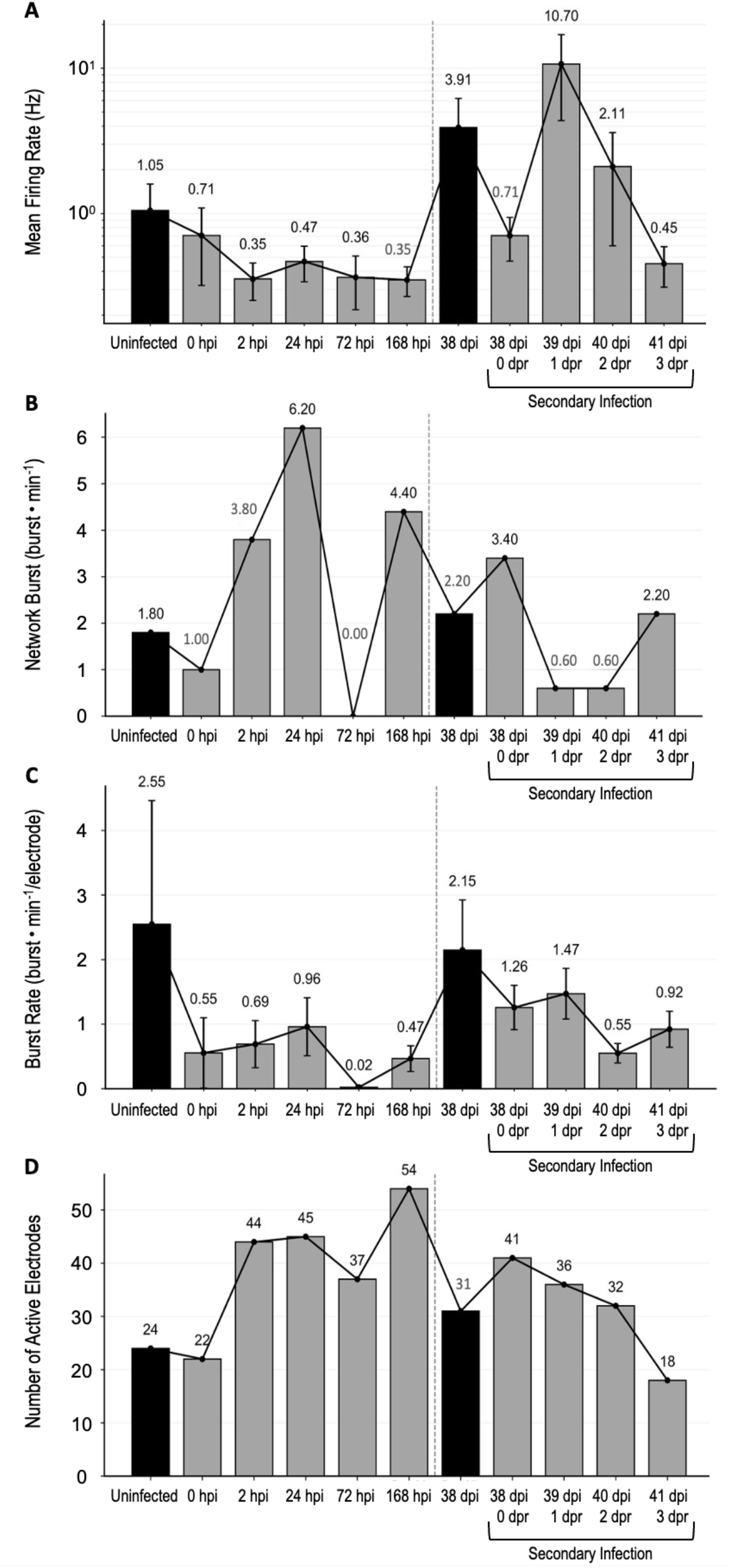
NGN2-derived forebrain neuron electrophysiology upon HHV-6A infection. MEA MEA recordings were used to assess electrophysiological activity in NGN2-derived forebrain neuron cultures at 29 days in culture and again upon a secondary infection (i.e., re-infection) at 67 days in culture both before and after HHV-6A infection. **(A)** Mean firing rate (MFR) across active electrodes. **(B)** Median inter-spike interval (ISI). **(C)** Burst rate per active electrode (BRE). **(D)** Number of active electrodes. Bars show summary values across active electrodes with error bars representing variability across active electrodes where applicable; the black line connects time points to show the longitudinal trend. Black bars indicate the uninfected baseline, and gray bars indicate HHV-6A infection time points.

For NGN2-derived neurons in culture for 29 days, MFR decreased quickly after primary infection (0-4 hpi) with a potential bump at 24 hpi (Figure 6, panel A, left field). Upon re-infection event at 38 days post-initial infection (67 days in culture), MFR immediately dropped, then increased at 24 hpi (1 dpr) from the second infection (Figure 6, panel A, right field) followed by a steady decline at 48 hpi (2 dpr) and 72 hpi (3 dpr), mirroring patterns observed in the NSC/NPC system. Upon initial infection median ISI showed a remarkable decrease at 24 hpi from the day of infection (0 hpi) providing some validation for the upward trend in MFR. Upon secondary infection, a modest decrease in ISI was observed (Figure 6, panel B, right field) at 24 hpi (1 dpr), which is supported by the concomitant increase in MFR (Figure 6, panels A). Burst rate during primary HHV-6A infection of NGN2 cultures shows a drop immediately after infection followed by steadily increasing trend in burst activity from 0-24 hpi followed by a sudden decline in burst activity at 72 hpi and, then a marked increase in bursting at 168 hpi (Figure 6, panel B, left field).

Upon secondary infection, the same pattern emerges with decrease in burst rate within hours of re-infection followed by an increasing trend at 24 hpi. However, at 48 hpi (2dpr), there was a significant drop in bursting with recovery by 72 hpi (3 dpr), supporting the notion that real changes in burst activity are occurring after viral infection (Figure 6, panel C). The number of active electrodes during the course of the trials is relatively stable from 2-72 hpi with a remarkable increase in the number of active electrodes at 168 hpi for primary infection (Figure 6, panel D, left field). While upon re-infection, there was a slight increase in the number of active electrodes (31 to 41) and then a modest decline (41 to 32) over 48 hpi (0-2 dpr) followed by a noted drop in the number of active electrodes at 3 dpr (Figure 6, panel D, right field).

### Virus Titers during the Course of Infection monitored by qPCR

Through all trials described above, periodic assessment of HHV6 titer was monitored to ensure that cultures were involved in productive infections. Specifically, virus-specific primers were used in conjunction with quantitative polymerase chain reaction (qPCR). Data show that virus titers increase over the course of infection, thus ensuring that recordings were taken. An example dataset shows increases in HHV6 titers from 24-72 hpi compared to uninfected culture controls (Figure S1).

## DISCUSSION

This protocol provides a detailed description, including a workflow diagram, demonstrating the utility of combining multi-electrode array (MEA) electrophysiology, immunofluorescence (IF), and molecular biology methods to understand virus-host dynamics in a neurotropic virus system. This workflow may also incorporate -omics approaches (e.g., RNAseq) and protein biochemistry (e.g., gel electrophoresis and western blot analysis) as explained below. However, for this report, we used immunofluorescence microscopy and MEA data to demonstrate the electrophysiological impacts of roseolovirus (i.e., HHV-6A) infection on neuronal activity in two distinct culture systems and in both 2D monolayer and 3D hydrogel formats. We used qPCR to titer virus working stocks and examine host susceptibility/viral productivity at different time-points post-infection.

A common critique when employing *in vitro* approaches to understand nerve cell responses to neurotropic agents (e.g., neuroactive compounds, viruses) is that responses observed in culture do not represent what may occur *in vivo*. Although this problem cannot be completely resolved, using multiple types of *in vitro* culture preparations provides some confidence that cellular responses are *bona fide* and will map to the in *vivo* condition, especially if responses in distinct preparations are consistent. In this study, we tested two distinct nerve cell culture preparations: (a) NSC/NPC-derived neuron-astrocyte mixed cultures; and, (b) NGN2-derived neuronal cultures. To ensure that differentiated cells from these cultures have matured to a state of electrogenicity, microscopy and pharmacology were employed. Brightfield microscopy confirms robust soma and neurite extensions/networks ∼50^+^ days after culturing in NSC/NPC-derived neuron-astrocyte cultures (see Figure 2, panels E and F) and ∼60^+^ days after culturing NGN2-derived neuronal cultures (see Figure 3, panels D and E). Both 2D and 3D BMM platforms may be seeded as describe in the Methods section. Upon applying two GABA_A_ receptor antagonists -bicuculine (Figure 1, panel A-D) and gabazine (Figure 1, panels E-H) – changes in MFR, ISI, and burst rate (BR) are evident. Specifically, inhibition of this inhibitory receptor system results in system excitation featuring increased MFR, decreased ISI, and/or increased BR. Other neuroactive compounds may be used. For example, in this report, we also applied nicotine, a potent cholinergic agonist. Although observations of changes in MFR were inconclusive, there was a drop in ISI and an increase in BR. It has previously been reported by our lab and others that during the differentiation process, glutamatergic and GABAergic cells seem to emerge prior to cholinergic and catecholaminergic neurotransmitter chemotypes. Thus, a robust GABAergic response was expected while an equivalent cholinergic response was not.

As an alternative way of determining what neuronal neurotransmitter chemotypes may be present in cultures, immunofluorescence using a fluorescent antibody-antigen system may be employed targeting antigens that are specific for certain neuron types. In this report, we not only demonstrated the presence of neurons (Figure 2, panel A and Figure 3, panel A) by targeting βIII-tubulin (i.e., Tuj1) as a neuron-specific antigen but also showed that the fluorescent signal for Tuj1 overlapped with HHV-6 gp60/110 fluorescence, confirming productive infection of neurons by HHV-6A. To identify the neuronal neurotransmitter chemotypes that may be serving as susceptible and permissive neuronal hosts, we targeted antigens vGluT1 and TH.

For both NSC/NPC-derived neuron-astrocyte and NGN2-derived neuronal cultures, a robust overlap between HHV-6A gp60/110 antibody fluorescence and vGluT1 signal was observed (Figure 2, panel B and Figure 3, panel B). Since vGluT1 is a vesicular glutamate transporter, it serves as a common marker for identifying glutamatergic cells. The ability for HHV-6A to productively infect glutamatergic neurons as shown in this report is support by prior work from our lab.^13^ For both NSC/NPC-derived neuron-astrocyte and NGN2-derived neuronal cultures some modest overlap was also observed between gp60/110 on TH and, separately, on vGluT1 signals (Figure 2, panel C and Figure 3, panel C, respectively). Since tyrosine hydrolase is a key enzyme in catecholamine production, it serves as a robust marker for catecholaminergic cells (e.g., dopaminergic neurons). The sparse overlap between HHV-6A gp60/110 and TH fluorescent signals raises the question of whether catecholaminergic cells are less susceptible and permissive to HHV-6A. Prior work from our lab in H9-derived neurons suggests that dopaminergic cells indeed may be susceptible.^13^ However, the relative susceptibility between dopaminergic and glutaminergic neurons remains under-studied.

Since the NSC/NPC-derived cultures contained glial cells, a GFAP fluorescent antibody was used to determine if glia (i.e., astrocytes) serve as productive hosts during HHV-6A infection. Indeed, a clear and robust overlap between HHV-6A gp60/110 and GFAP fluorescent signals emerged. This, of course, suggests that glial cells are also susceptible to HHV-6A infection, which is consistent with prior work by our lab and others on the ability of HHV-6 to infect glia.^5,7,13^

Other markers may be used for IF assays to determine what neuronal neurotransmitter chemotypes are present within cultures and susceptible to viral infection. For example, markers for cholinergic neurons such as Choline Acetyltransferase (ChAT) or Acetylcholine esterase (AChE) or subunits for nicotinic acetylcholine receptors (nACHRs) may also be employed. Integration of data from immunofluorescence experiments and MEA recordings using pharmacological agents and/or viral infection (Figure 1) can address a myriad complex research questions regarding neurotropic virus infection. This may include discerning the relative contribution of distinct neuronal neurotransmitter chemotypes to neural network signaling perturbations during viral infection.

Ultimately, the methods presented here are focused on understanding the role of virus infection (i.e., HHV-6A infection) on nerve cell function. To this end, MEA recordings were taken from NSC/NPC-derived and NGN2-derived cultures both immediately prior to and for several days post-infection (see Figures 4 and 5). Without any neuroactive compounds applied, changes in electrical activity within these cultures were observed upon infection with HHV-6A at MOI=1. In some readily observable changes in MFR, ISI, or BR could be observed between consecutive time-points (e.g., from 24-48 hpi). However, this was not typical for many recordings. Although responses between consecutive time-points may not show any remarkable change in MFR, ISI, or BR, for many responses, clearly observable changes are observed between earlier and later timepoints in the series (e.g., between 24 hpi and 168 hpi), suggesting that the trends observed between consecutive timepoints represent *bona fide* electrophysiologic patterns.

A common issue in using nerve cells derived from iPSCs for in vitro preparations is time-in-culture. Different preparations, cell lines, differentiation protocols, and other variations have unique time courses of cellular maturation. Not only does this impact the emergence and maturity of electrogenic activity but also susceptibility profiles to viral infection. As demonstrated in this report with the NGN2-derived neuronal cultures (see Figure 5), the workflow presented is amenable to testing cellular responsiveness to viral infection at multiple time points after seeding the culture and accommodates questions regarding changes in nerve cell responsiveness during the maturation process.

Although it is beyond the scope of this report, it is important to acknowledge that the culturing, electrophysiology, immunofluorescence, and pharmacology techniques presented here may also be coupled with biochemistry and multi-omics approaches. As a tacit example, we demonstrate that qPCR (molecular biology) may be employed to measure virus titer during the course of infection (see Figure S1). Our lab also applies techniques in protein gel electrophoresis such as SDS-PAGE and Native PAGE and western blot analysis (protein biochemistry) as well as RNAseq (transcriptomics) to obtain additional data sets that may support or refute working hypotheses. For example, we have tested the hypothesis that nerve cell susceptibility to HHV-6 infection in differentiated cell lines may be a function of receptor expression. HHV-6A preferentially uses the Cluster of Differentiation 46 (CD46) cell surface protein for attachment and entry, while HHV-6B preferentially uses CD134. Employing methods in immunofluorescence, gel electrophoresis, RTqPCR, we were able to address this hypothesis.

## CONCLUSION

This report introduces a well-defined multi-method workflow to integrate data from MEA recordings, immunofluorescence, molecular biology, biochemistry, and multi-omics approaches in differentiated human iPSCs to elucidate the impacts of viral infection on nerve cells. Specifically, we provide detailed methods on how to culture NSC/NPC-derived neuron-astrocyte cultures and NGN2-derived neuronal cultures in both 2D monolayer and 3D “hydrogel” formats. These cultures are then used to characterize the impacts of roseolovirus (i.e., HHV-6A) infection on nerve cells electrophysiological responses. IF and biochemical data are used in conjunction with MEA data to consider complex questions regarding cell tropism and host relative susceptibility to HHV-6A infection. The workflow presented (see Figure 1) may be expanded to integrate data from other techniques to achieve a more comprehensive understanding of virus-host dynamics in neural systems.

## ACKNOWLEDGMENTS

This work was support by a U.S. National Science Foundation, Division of Biological Sciences, Biology Integration Institute grant (Award No. 2119968; PI/PD-Ceballos).

---

**Figure S1:**
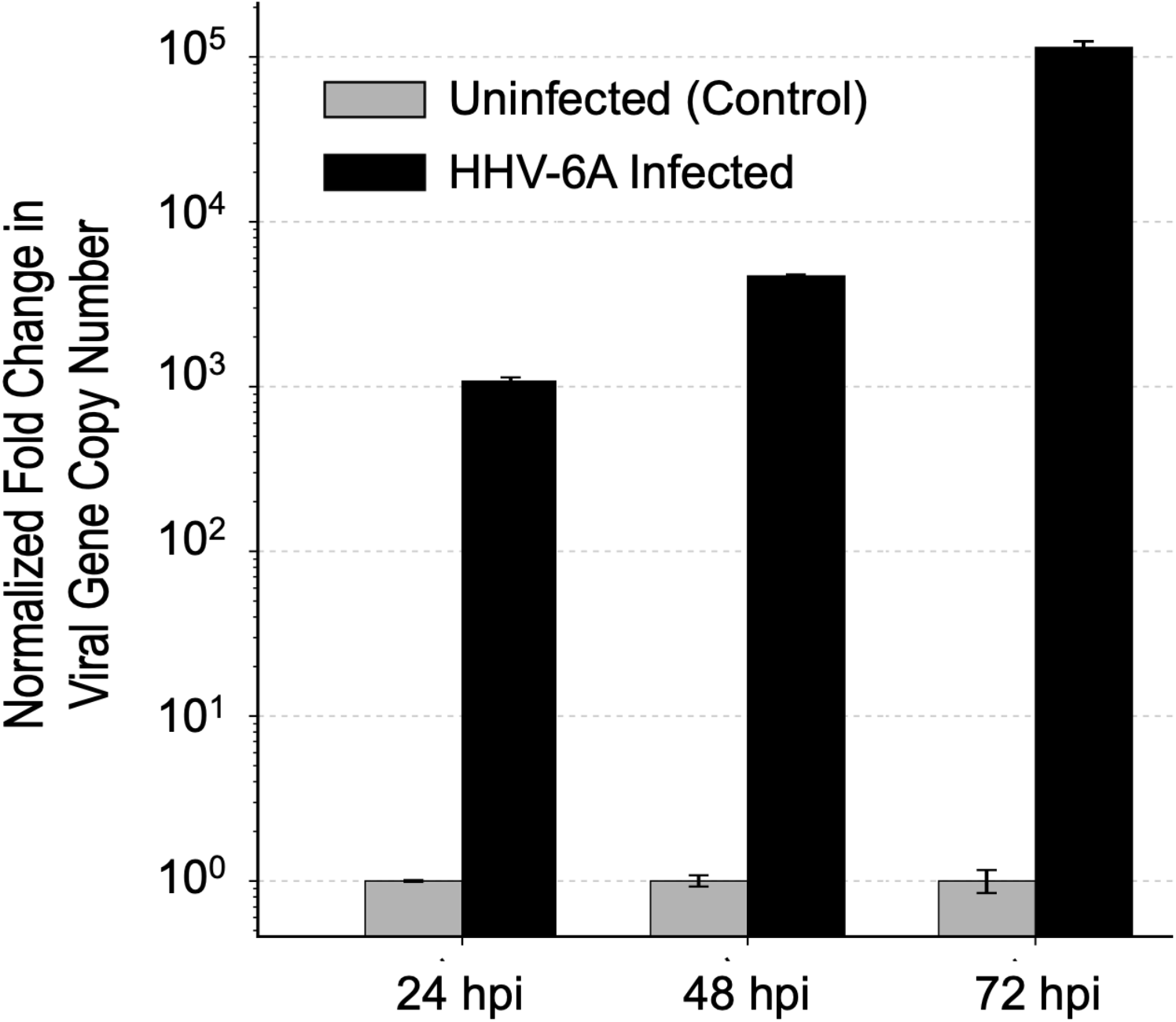
HHV-6A viral gene counts after infection over select time-points post-infection. Normalized fold change in HHV-6A viral gene copy number in HHV-6A infected nerve cell cultures at 24, 48, and 72 h post-infection (black bars) versus uninfected controls (grey bars).

